# Serial femtosecond and serial synchrotron crystallography yield data of equivalent quality: a systematic comparison

**DOI:** 10.1101/2020.08.21.257170

**Authors:** P. Mehrabi, R. Bücker, G. Bourenkov, H.M. Ginn, D. von Stetten, H.M. Müller-Werkmeister, A. Kuo, T. Morizumi, B.T. Eger, W.-L. Ou, S. Oghbaey, A. Sarracini, J.E. Besaw, O. Paré-Labrosse, S. Meier, H. Schikora, F. Tellkamp, A. Marx, D.A. Sherrell, D. Axford, R. Owen, O.P. Ernst, E.F. Pai, E.C. Schulz, R.J.D. Miller

**Author notes:** These authors contributed equally to this work (P. Mehrabi, E.C. Schulz).

## Abstract

For the two proteins myoglobin (MB) and fluoroacetate dehalogenase (FAcD), we present a systematic comparison of crystallographic diffraction data collected by serial femtosecond (SFX) and serial synchrotron crystallography (SSX). To maximize comparability, we used the same batch of crystals, the same sample delivery device, as well as the same data analysis software. Overall figures of merit indicate that the data of both radiation sources are of equivalent quality. For both proteins reasonable data statistics can be obtained with approximately 5000 room temperature diffraction images irrespective of the radiation source. The direct comparability of SSX and SFX data indicates that diffraction quality is rather linked to the properties of the crystals than to the radiation source. Time-resolved experiments can therefore be conducted at the source that best matches the desired time-resolution.

## Introduction

Structural biology has been highly successful in providing three-dimensional information about the architecture of biomolecules. Usually these structures display equilibrium state conformations, important snapshots that provide insight into functional states of these biomolecules^1^. The advent of X-ray free-electron laser (XFEL) sources promised the possibility to study ultra-fast timescales down to the femtosecond domain to watch conformational changes and even chemical reactions^2^. Addressing irreversible reactions at these ultra-fast timescales and guaranteeing homogeneous reaction initiation required the use of micron-sized protein crystals. However, the extremely intense beam-brightness of XFEL sources will destroy those protein micro-crystals during single exposures. This dilemma was circumvented by serial data collection schemes in which hundreds of thousands of micro-crystals are exploited to each provide a single still diffraction pattern. Due to the *diffraction-before-destruction* approach, which allows the collection of a useful diffraction pattern by outrunning the onset of damage, it has been argued that serial femtosecond crystallography (SFX) data from XFELs are *de-facto* radiation damage free ^3–5^.

Soon after the first protein crystal structures were solved from SFX data, the method was adapted for use at synchrotrons giving rise to serial synchrotron crystallography (SSX)^6,7^. The majority of SSX experiments thus far involved static structures. However, since most enzymes have median turnover times in the hundreds of milliseconds range, synchrotrons represent a valid alternative for time-resolved experiments. Historically exploited via Laue-diffraction on single-crystals ^8,9^, recently this has been the subject of serial data collection approaches to study enzymes on timescales from milliseconds to many seconds^10–15^. The results of those studies have sparked renewed and expanded interest in serial crystallography methods, and the implementation of time-resolved SSX at a number of synchrotron beamlines can be expected, in particular as improving synchrotron technology will also bring faster time-domains into reach.

Due to the large number of photons compressed into the X-ray laser pulses SFX exposure times can be very short, down to the femtosecond domain^2^. As a consequence of the more limited photon flux at synchrotron sources accordingly longer exposure times are needed in SSX, and consequently radiation damage cannot be fully outrun. This is particularly severe for time-resolved experiments because data need to be collected at room-temperature leading to much increased radiation sensitivity. However, while radiation damage is more pronounced at synchrotron sources, much of its deleterious effects can be mitigated by using low-dose exposure, which still leaves the electron density interpretable with little ambiguity ^16,17^. The dose causing a 50% drop in overall diffraction intensity for lysozyme at room temperature SSX has been reported to be 380 kGy ^18^. While this represents the upper dose limit for serial crystallography experiments, which should not be exceeded at room temperature, the onset of site-specific damage could be observed at a much lower dose of approximately 80 kGy ^18^. Although this value is much lower than doses encountered during typical cryo experiments, at modern microfocus beamlines it is still possible to collect multiple frames from a single micro-crystal at room temperature within this dose limit ^17–19^.

For a systematic comparison of SFX and SSX data, we analyzed two different proteins at both radiation sources. To minimize sources of potential discrepancies, we used the same batch of micron-sized crystals, the same sample delivery device, and the same data analysis software for each of the data sets.

## Results and discussion

To evaluate potential differences in quality between crystal diffraction data obtained from XFEL (SFX) and those from synchrotron sources (SSX), we exploited two well established protein systems: i) the relatively radiation tolerant fluoroacetate dehalogenase (FAcD) and ii) the highly radiation sensitive myoglobin (MB). FAcD is a homodimeric protein from the soil-bacterium *Rhodopseudomonas palustris* and one of the very few enzymes that can cleave fluorine carbon bonds. FAcD shows half-of-the-sites reactivity and incorporates a covalent substrate-enzyme intermediate into its SN2 substitution mechanism ^15,20,21^. MB is a monomeric heme protein involved in the oxygen exchange of mammalian muscle tissue. It was the first protein that had its three-dimensional structure determined and is known to be highly radiation sensitive due to the iron center of its heme chromophore^22,23^.

For sample delivery, we made use of our previously described fixed target approach^16,24–27^. Briefly, approximately 25,000 protein micro-crystals are mounted on a silicon support chip in random orientations. These chips are then raster-scanned in the X-ray beam by means of a fast and accurate closed-loop piezo translation stage system. This system allows the collection of data from up to 4 chips (i.e., approximately 100,000 microcrystals) per hour. The modular architecture of this system enables its use at different synchrotron and XFEL end-stations ^14,16,19,24–28^. For the present comparison, data were collected at SACLA at the RIKEN SPRING-8 center and the EMBL beamline P14 at the PETRA-III synchrotron at DESY. Importantly, in addition to applying the same data-collection hardware we used protein crystals from the same crystallization batches in both experiments. The micro-crystals of both proteins were isomorphous and of the same physical dimensions. This should reduce any crystal quality differences that may influence overall data quality parameters. Moreover, to reduce any further mismatches in the analysis, the data were processed with the same software package, CrystFEL 0.8.0 ^29^.

### Global data quality comparison

The global data quality parameters for both proteins indicate close equivalence of the SSX and SFX data over the whole resolution range (**Table I, Figure 1**). For both proteins, signal to noise ratio (SNR), multiplicity, R_split_, and overall completeness are almost identical for SFX and SSX data (**Figure 2**). Only minor resolution differences can be observed, as indicated by the resolution-shell half-set correlation coefficients (CC_1/2_), R_split_, and refinement *R*_free_ values. While the MB dataset shows virtually no differences in B-factors between the SSX and SFX dataset, the B-factors are slightly lower for the SFX dataset of FAcD (**Table I**). This may correlate with the marginally better data statistics in the high-resolution shell of FAcD as made evident by SNR and R_split_ values. It is conceivable that the FAcD crystals would have diffracted to a slightly higher resolution in the XFEL beam relative to the synchrotron beam as indicated by the B-factors. The current resolution cutoff was defined by the detector distance at the XFEL. In addition to the integration statistics, the general quality of the refined structures needs to be assessed. Globally this can be achieved by comparing the free *R*-values (*R*_free_) of the SSX and SFX structures (**Table I**). The absolute percentage differences between the overall *R*_free_ values for FAcD and MB are negligible, while for the high-resolution shell absolute *R*_free_ value differences of ∼4.6 % and ∼1.8 % can be observed, respectively. On a per-residue basis, model differences between the SFX and SSX structures can conveniently be assessed via the root-mean-square deviation (r.m.s.d.) and *B*-factor values. The all-atom r.m.s.d. of the SFX and SSX structures for FAcD is 0.12 Å, compared to a coordinate error of 0.20 Å. For MB, the all-atom r.m.s.d. of the SFX and SSX structures is 0.13 Å, compared to a coordinate error of 0.23 Å.

**Table 1:**
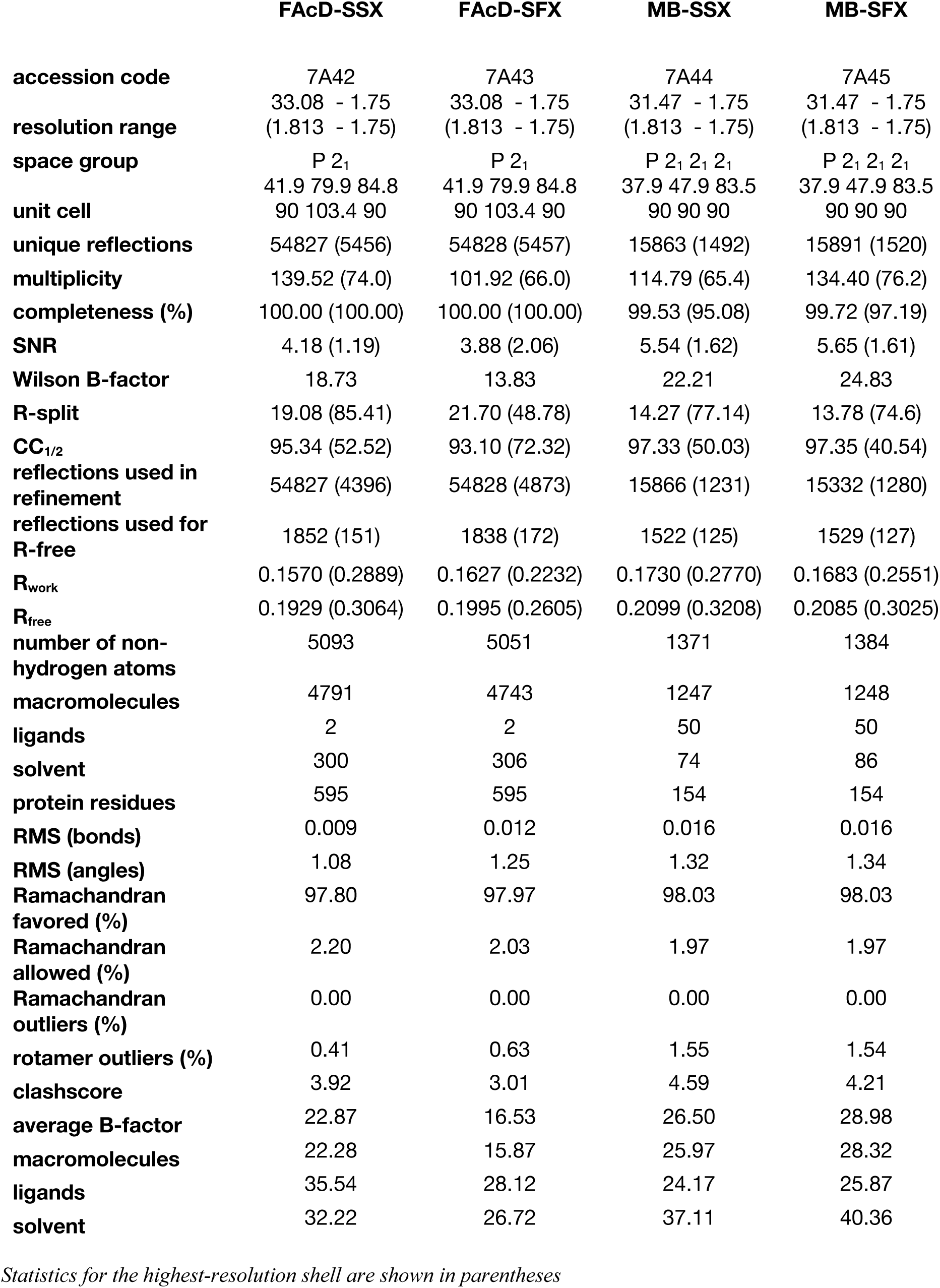
Data collection and refinement statistics.

**Figure 1:**
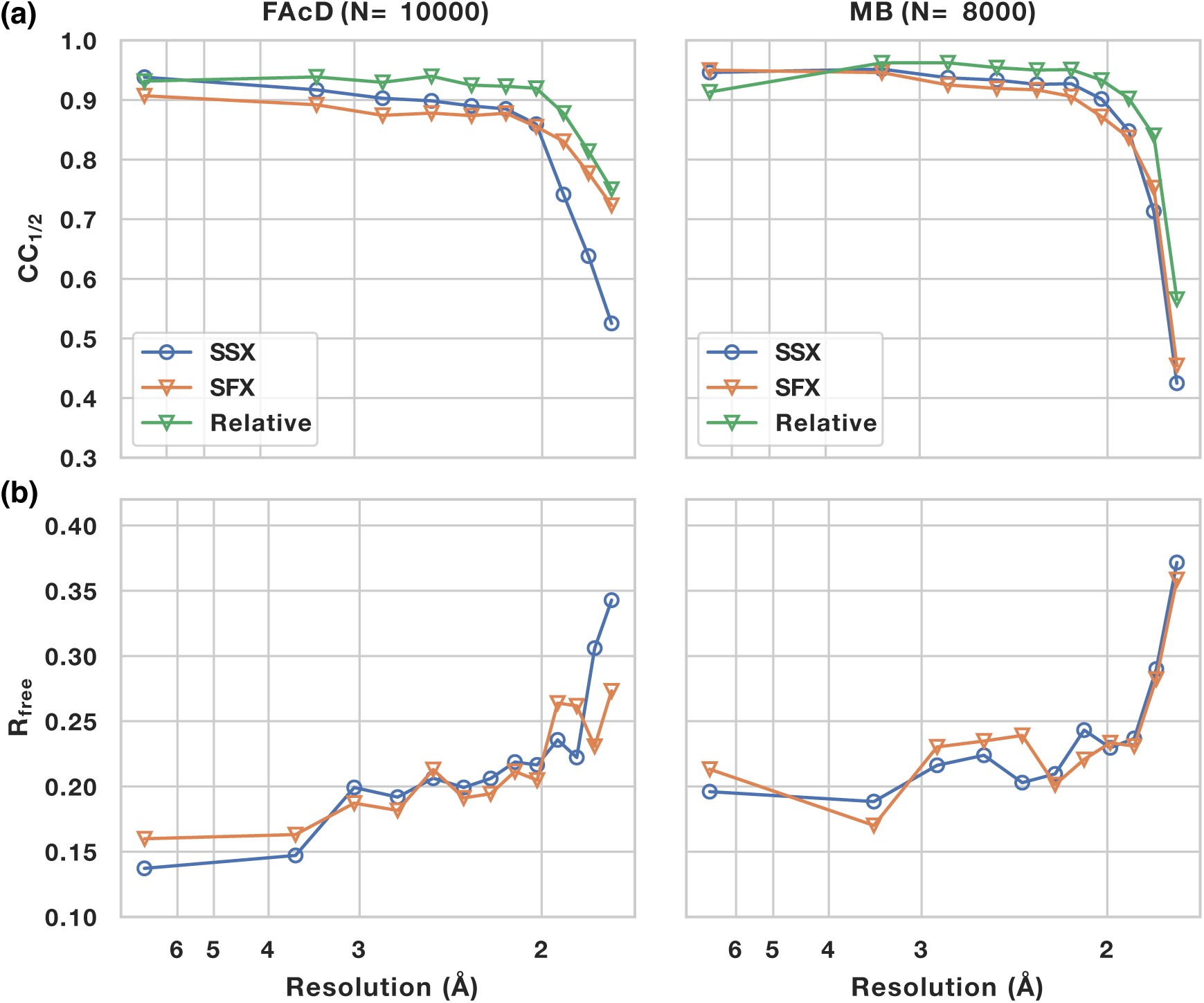
Global data quality parameters as function of resolution. (a) half-set correlation coefficient for SSX (blue, open circles) and SFX (orange, open triangles) for FAcD and Myoglobin (MB) data sets. The green curve displays the relative correlation coefficient between SFX and SSX data. The datasets were limited to the same number of diffraction images (10,000 for FAcD and 8,000 for myoglobin) and the same resolution cutoff was applied to SSX and SFX data. (b) corresponding refinement R_free_ values.

**Figure 2:**
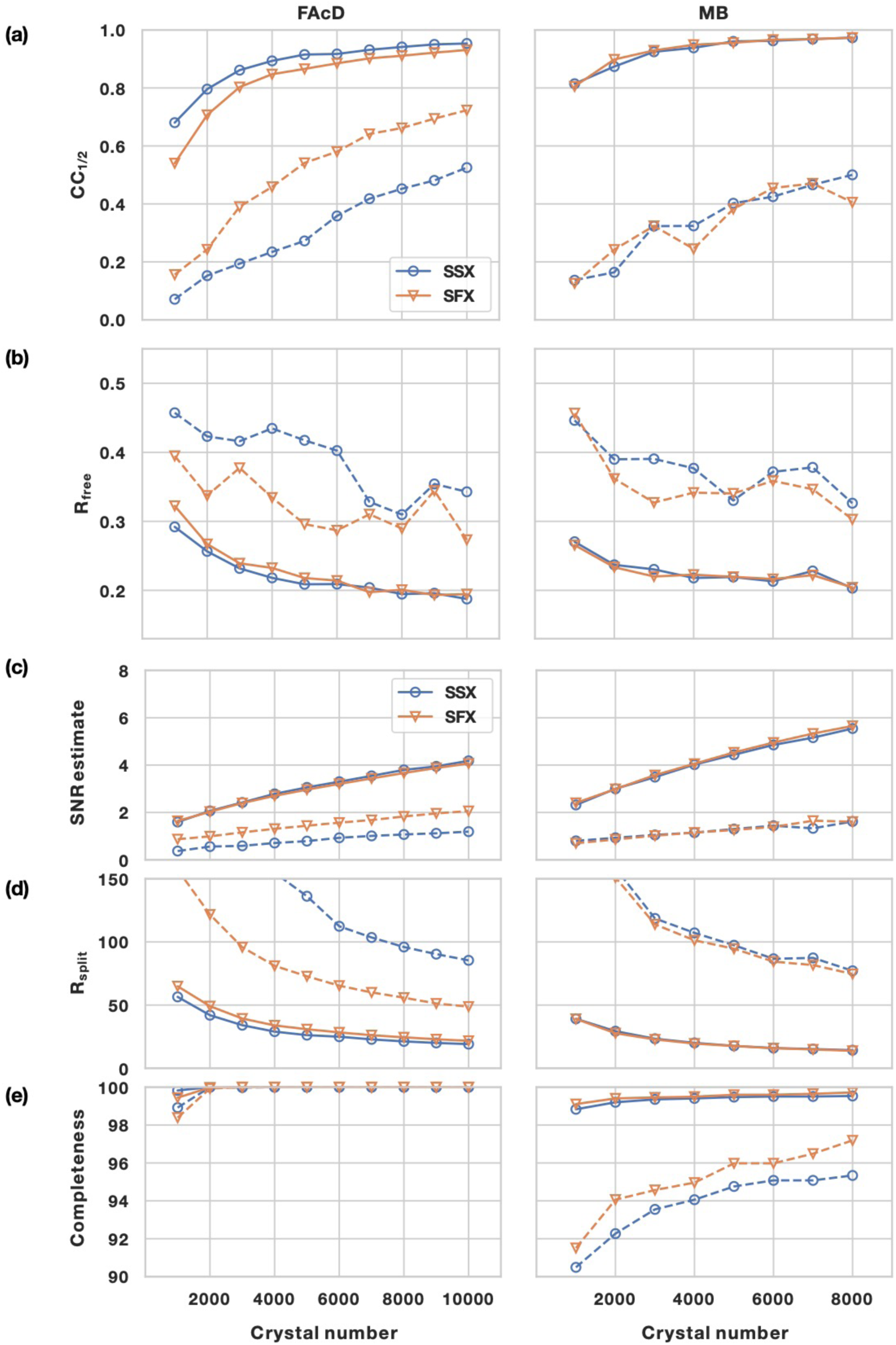
Global data quality parameters as function of data set size (number of crystals). Global data quality parameters for SSX and XFEL data sets: (a) CC_1/2_ values, (b) R_free_ values, (c) SNR estimate, (d) R_split_ values, and (d) completeness. SSX data are displayed as blue, open circles, SFX data as orange, open triangles. Overall values are shown as solid lines, highest resolution shell values as dashed lines.

By collecting room temperature data from the same batch of micron-sized crystals, using the same sample delivery device, as well as the same software for data-analysis, we have attempted to maximize comparability of data collection at XFEL and synchrotron sources. Indeed, this experimental equivalence is reflected in very similar data statistics for the SFX and SSX data for both protein systems. The data quality indicators obtained (such as CC_1/2_, R_free_ and r.m.s.d.) are nearly identical for both radiation sources. However, thus far this statement only holds true for crystals of suitable size (i.e. on the order of a few micrometers), that enable data collection at synchrotrons. Clearly XFELs can make use of much higher flux densities, thereby accommodating much smaller crystals, down to the nanometer scale, which currently is outside the range of synchrotrons^30^. In summary, by normalizing all experimental parameters, this comparison demonstrates that in (static) serial X-ray crystallography data quality is predominantly a crystal-dependent property. Our observations reveal that neither obtainable resolution nor CC_1/2_ or R_free_-values strongly depend on the radiation source. As a consequence, the diffraction properties of protein microcrystals can be explored at synchrotron sources to test their suitability for XFEL experiments. This would relieve XFEL beamlines of the high burden of screening time as well as increase opportunities for testing crystals as beamtime at synchrotrons is much more readily available. This not only includes static data collection, but of course also extends to time-resolved experiments, where crystals can be used for experiments at synchrotron sources for observing somewhat slower time domains.

### Scaling with data set size

Subsequently, we correlated the quality of the datasets with the number of crystals used for structure determination, randomly picking sub-sets of the total acquired datasets. Analysis of correlation coefficients, R_free_ values, SNR, R_split_, and completeness again shows a high degree of similarity between SSX and SFX data. Although the SFX data were recorded with a bandwidth approximately 2 orders of magnitude larger than available at monochromatic synchrotron beams, the number of diffraction patterns needed to converge is similar for both datasets. While true Laue serial diffraction data require fewer images for convergence, this criterion is apparently not met by the increased bandwidth of XFEL beams compared to monochromatic synchrotron beams^31^.

In agreement with our previous observations, the datasets of both proteins show almost full completeness, that is over 95% in the highest resolution shell, even for a low number (< 1000) of diffraction patterns^32^. This indicates that for serial data collection, completeness should be the first data quality metric to be assessed before turning to other quality indicators such as CC_1/2_ or *R*_split_. We argue that if the data are not fully complete other data-quality indicators are probably meaningless. For both FAcD and MB, the overall quality indicators CC_1/2_ and *R*_free_ start to converge to reasonable levels at approximately 5,000 diffraction images, by standard assessments (**Figure 2**). While there are only minor improvements for overall data quality indicators, the high resolution-shell quality indicators still undergo substantial improvements if more than 5,000 diffraction patterns are included. The FAcD and MB crystals used in this analysis were of monoclinic and orthorhombic symmetry, respectively – representing the two most commonly encountered crystal systems in the PDB. This suggests that for the vast majority of cases reasonable data statistics should be achievable with a similar number of diffraction images. For triclinic symmetry (representing less than 5% of the structures in the PDB), however, a higher number of diffraction patterns would be required.

This corroborates our previous findings that protein structures and even protein small-molecule complexes obtained by SFX and SSX can be solved from ∼5,000 diffraction patterns - or fewer if high symmetry permits - yielding reasonable electron density ^32,33^. However, merging and refinement statistics, as well as the quality of electron density, clearly improve if further diffraction patterns are added. This can be especially important in time-resolved analyses to reveal low occupancy details or to distinguish subtle electron density differences from the noise.

Thus, as a rule of thumb, the present data suggest to obtain approximately 5,000 diffraction patterns, as at this point the overall dataset quality has mostly to converged to reasonable quality indicators. Provided micron-sized crystals are used, SFX and SSX appear to be equivalent alternatives not only regarding the quality of the data but also with respect to the required number of diffraction patterns. Exciting developments in nano-focus X-ray beamlines combined with the increased brilliance of next generation synchrotron sources may narrow the gap of useful crystal sizes in the future, thereby also enabling the use of nano-crystals at synchrotrons ^34,35^. Since microsecond time-resolution can also be obtained at current generation synchrotron sources, experiments on XFELs can therefore be focused on ultrafast time-resolution experiments, the time domain on which XFELs are essential.

### Radiation damage

For traditional crystallographic experiments at synchrotrons, radiation damage is a key limiting factor in obtaining high resolution structures even at cryogenic temperatures. Damage is caused by the interaction of the X-ray photons with the electrons in the protein crystals and the energy they deposit in the sample, either caused by the photoelectric effect or inelastic scattering (Compton scattering) of X-rays leading to primary, secondary, and tertiary damage ^36,37^. The traditional Garman limit for cryo-temperature structures is 30 MGy^38^. The situation is worse at room-temperature, typically encountered during serial crystallography experiments. Radiation damage manifests either globally as a loss of diffraction intensity, or specifically via chemical modification of protein residues, i.e. breaking of disulfide bonds^36,39^. Recently, dose limits for room-temperature SSX were inferred from lysozyme crystals, indicating a half-diffraction dose of 380 kGy and site specific damage starting to appear at approximately 80 kGy ^18,^. Notably, the authors implied that the onset of site-specific damage may occur at much lower doses for radiation sensitive proteins. This agrees well with previous findings that the dose-limit at room temperature is highly sample-dependent varying over an order of magnitude for 15 different structures^40^. In comparing synchrotron and XFEL sources radiation, one needs to further distinguish between direct and indirect damage. While direct damage occurs via the primary X-ray absorption at an atom in the protein, indirect damage occurs via X-ray absorption in the bulk solvent ^36^. The latter leading to the formation of reactive oxygen species on the ns-ms timescale that can diffuse through the crystal^41^. Thus, the advantages of XFELs with respect to radiation damage originate in their short pulse lengths, which outrun those damage mechanisms caused by the diffusion of radicals encountered during longer exposure times at synchrotrons ^42,43^. Due to the short pulse length (10 fs) and high intensity of the SACLA beam, established dose criteria cannot be applied to SFX data in an unmodified manner. For reference, we therefore used an updated version of RADDOSE (RADDOSE-XFEL), which now also offers to correctly calculate XFEL as well as SSX doses. We estimated the average dose in the exposed region, indicating that the 80 kGy dose limit was exceeded by a factor of ∼2 for the SSX data (**Table II**)^44^.

**Table II:** Results from RADDOSE-XFEL.

|  | average diffraction<br>weighted dose (MGy) | RADDOSE-XFEL<br>average dose exposed<br>region (MGy) | RADDOSE-3D style<br>average dose exposed<br>region (MGy) |
| --- | --- | --- | --- |
| **MB-SSX** | 0.252 | - | 0.184 |
| **MB-SFX** | - | 0.200 | 1.022 |
| **FAcD-SSX** | 0.229 | - | 0.167 |
| **FAcD-SFX** | - | 0.163 | 0.959 |

No global structural differences were observed in either FAcD or MB. To address the site-specific differences between the SFX and SSX data we utilized a resolution-shell scaling tool to obtain isomorphous differences densities^45^ (*Ginn et al. – under review*). The obtained difference density maps do not display the absolute damage but rather indicate relative differences between the two datasets (**Figure 3**). For FAcD the difference density peaks are evenly distributed over the whole structure, indicating no preferred site of radiation damage. By contrast for MB, damage is less homogenously dispersed throughout the structure; rather local differences are primarily concentrated around certain areas such as the radiation sensitive heme-centre. This agrees with canonical radiation damage mechanisms induced by the reduction of the iron centre ^23^ (**Figure 3**). These findings further support the notion that SFX and SSX data are largely comparable and no major differences with respect to radiation damage can be observed. While SSX data are clearly not radiation damage free, the serial data collection approach apparently allows to collect data that show only minimal differences to SFX-data, which outrun the most deleterious radiation damage effects – even for very radiation damage-sensitive proteins such as myoglobin. This implies that only in very critical cases radiation damage is a crucial criterion for the choice of the radiation source. While the present SSX data were collected at doses of approximately 170 kGy, with new rapid acquisition detectors (Eiger) it is possible to collect SSX data with < 2 ms exposure time thus at much lower doses (< 5kGy), further mitigating the differences between SFX and SSX data^18^.

**Figure 3:**
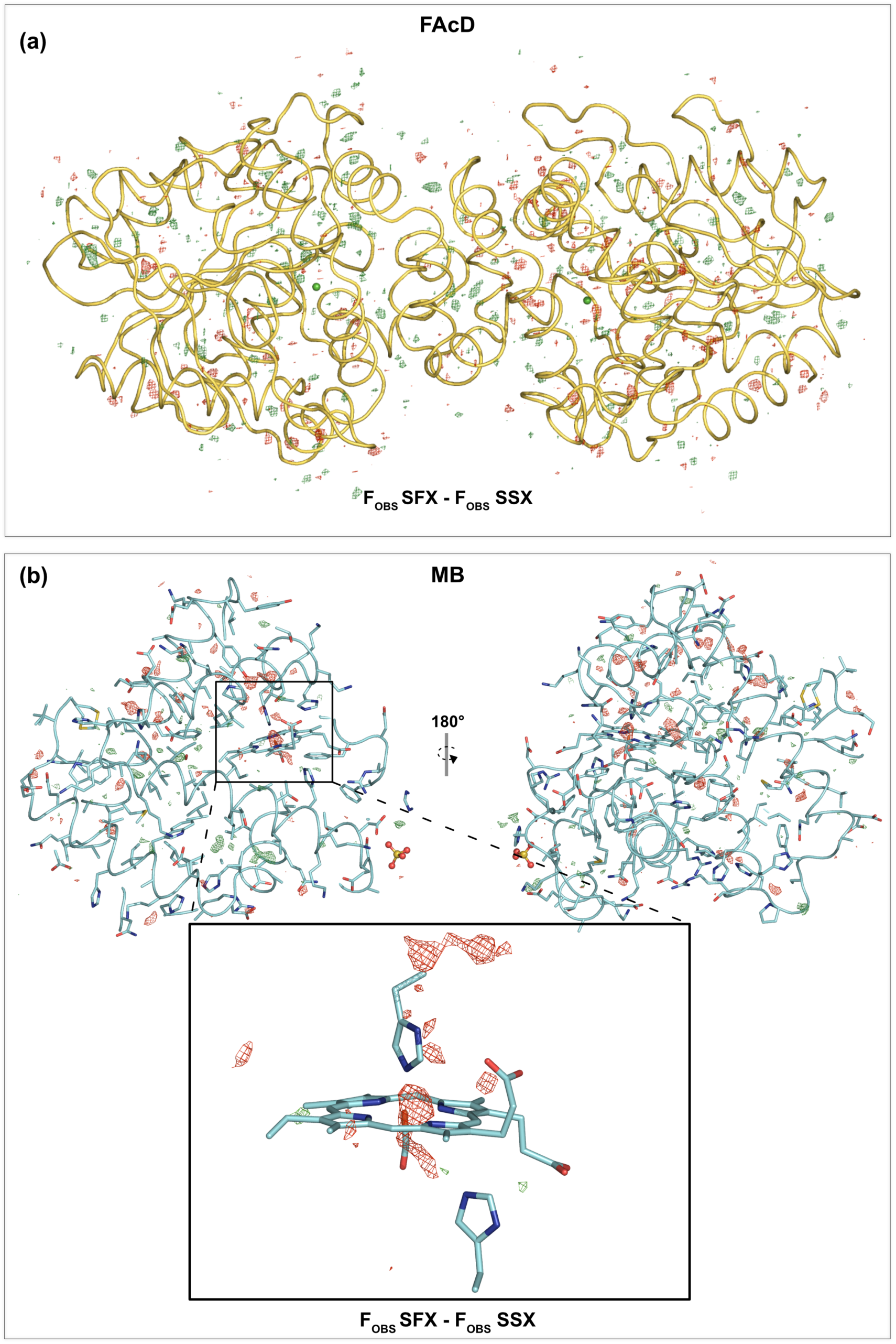
Isomorphous difference maps for FAcD and MB, respectively. Resolution shell scaled F_OBS_ SFX - F_OBS_ SSX difference maps of FAcD (a) and MB (b). Difference map peaks are homogenously distributed over FAcd, while difference map peaks are more concentrated around the radiation-sensitive heme center in MB. Proteins are shown as cartoon representations, all maps are shown at +/- 3 sigma.

## Conclusions

Data obtained via serial crystallography approaches at synchrotron and XFEL sources can be of equivalent quality. Therefore, data quality is a crystal-dependent property and does not depend on the radiation source. We conclude that as a rule of thumb ∼5,000 diffraction patterns are required to obtain data statistics and electron density maps of reasonable quality; higher numbers may be required to tease out lowly populated electron density states. Further, our findings suggest that it is possible to tailor SSX data collection, to largely avoid the effects of radiation damage thereby preserving the biological interpretability of the results. Due to the overlapping data quality, the choice of the radiation source should primarily be guided by the required time resolution for time-resolved experiments.

## Methods

### Protein expression, purification and crystallization

Recombinant fluoroacetate dehalogenase (FAcD) was purified from *Escherichia coli* BL21(DE3) as described ^20^. FAcD was extracted from *E. coli* cell-free lysate using Ni-chromatography with subsequent cleavage of the His_6_-tag using TEV protease. Size exclusion chromatography was completed using 50 mM Tris-H_2_SO_4,_ pH 8.5 and 150 mM NaCl and buffer exchanged to remove the NaCl as a final step in purification. FAcD crystals were grown in crystallization buffer (18-20 % (w/v) PEG3350, 200 mM CaCl_2_, and 100 mM Tris-HCl, pH 8.5). From these crystals, a microseed stock was generated at a 4-8% higher PEG3350 concentration, using a seed bead kit from Hampton research (HR2-320). Microcrystals were produced using batch crystallization; 100-200 µl of seed stock and an equal volume of 0.5 mM FAcD solution were mixed. Within 24-72 hours crystals grew to approximately 20 × 20 x 10 µm^3^ in size. CO-bound sperm whale myoglobin was prepared and crystallized as described previously ^16,25^.

### Serial synchrotron crystallography data collection

Single crystalline silicon chips provide a suitable scaffold material for fixed target crystallography. They hold randomly oriented crystals in precisely defined, bottomless wells (also called features). 100-200 µl of a suspension of crystals were loaded onto a single chip by applying vacuum suction as described previously. Chip fabrication and sample loading process are described in detail in^25–27^. Serial synchrotron crystallographic (SSX) diffraction data were collected at room temperature (294 K) at EMBL beamline P14 at DESY, Hamburg, using our previously described fixed-target setup ^16,24^. Crystals were not rotated during an exposure; still images were recorded at a wavelength of 0.976 Å and an exposure time of 37 ms using an Eiger 16M detector (Dectris, Switzerland). SSX diffraction data were processed using CrystFEL 0.8.0 ^29^.

### Serial femtosecond crystallography data collection

Serial femtosecond crystallography (SFX) diffraction data of FAcD and MB were collected at room temperature (294 K) at SACLA at SPRING-8, Japan, using the same setup as described above. SFX diffraction data were processed using CrystFEL 0.8.0 ^29^.

### Molecular replacement and refinement

The structures were determined by molecular replacement in PHASER ^46^ using PDB IDs 5K3D and 5JOM as search models for FAcD and myoglobin, respectively. Structure refinement was completed by iterative cycles of refinement in *phenix*.*refine* ^47^ and manual model building in COOT ^48,49^.

### Dose and radiation damage

Doses for each dataset were calculated using the software RADDOSE-3D (v. 4.0) and its subprogram RADDOSE-XFEL, respectively ^44^. Cuboid crystal forms with dimensions of 20 × 20 x 10 µm^3^ for MB, and 20 × 20 x 15 µm^3^ for FAcD were applied. For the SSX data, a Gaussian beam profile with a flux of 4^12^ ph/s and a full width half maximum (FWHM) beam of 5 × 10 µm^2^, with a rectangular collimation of 15 × 30 µm^2^ was considered at an energy of 12.703 keV for an exposure time of 37 ms and a rotation wedge of 0°. RADDOSE-3D was run with a resolution of 5 pixels per micron and default angular resolution for a wedge size of 0°. For the XFEL data, a Gaussian beam profile, with an uncollimated FWHM size of 4.7 × 3.4 µm^2^ and an energy of 10.2 keV, a pulse energy of 0.4 mJ and an energy distribution of 0.5% (FWHM) as well as an exposure time of 10 fs was applied. RADDOSE-XFEL was run with a resolution of 0.5 pixels per micron and simulated 1,000,000 photons. Results from 3 runs were averaged and are summarized in **Table II**. Isomorphous difference maps were calculated in analogy to a previously published approach scaling the datasets to their resolution shells ^45^. The SFX datasets were scaled to the reference SSX datasets, by dividing the limiting resolution range into 20 equal volume bins containing similar numbers of reflections. The reflection amplitudes in each bin were multiplied by a constant scale factor, calculated to set the average amplitude in each bin equal to that of the reference. The code for shell-scaling is part of the Vagabond software suite (Ginn et al., unpublished). The resulting scaled datasets were plugged into sftools (CCP4 software suite) and difference structure factors were calculated F_obs-SFX_ – F_obs-SSX_ for both FAcD and MB respectively ^50^.

### Data analysis

Molecular images were generated in PyMOL (Schrödinger LLC).

## Author Contributions

E.C.S and P.M. initiated the project; P.M., T.M. and A.K. prepared the protein and crystals;. E.C.S, P.M., D.v.S, H.M.-W., A.M., H.S, H.G. D.S., D.A., R.O., F.T., A.K., J.E.B., T.M, W.-L.O., J.B., O.P.E., E.F.P., R.J.D.M. collected the data; E.C.S, P.M., R.B., H.G., B.T.E., and A.M. processed the data; E.C.S, P.M., R.B., D.v.S. and G.B. analyzed the data; E.C.S, P.M. and R.B. wrote the manuscript. All authors discussed and corrected the manuscript.

## Acknowledgements

The SSX data was collected at beamlines P14 operated by EMBL Hamburg at the PETRA III storage ring (DESY, Hamburg, Germany). We would like to thank Drs. A.R. Pearson (University of Hamburg), S. Horrell (Diamond Light Source), M. Agthe, and T.R. Schneider (EMBL Hamburg) for their assistance in using the beamline, and for their continuous support and helpful discussions in the implementation and improvement of SSX instruments at EMBL Hamburg. We acknowledge Dr. T. White for most helpful discussions on data processing. The XFEL experiments were performed at the BL3 of SACLA with the approval of the Japan Synchrotron Radiation Research Institute (JASRI) (Proposal No. 2016A8036 and 2016B8052). We thank the beamline scientists for their excellent support.

